# Synthesis and validation of a metal-organic framework for functional quenching of fluorescent DNA

**DOI:** 10.1101/2025.01.16.633169

**Authors:** James Goodrich, Ali Beladi, Konyin Joshua, Humza Bhatti

## Abstract

Access to practical research experience is typically limited for life science undergraduates at UK universities. This is confounded by a lack of time to pursue research given course commitments, and the competition for limited spaces on internships or other schemes. To overcome these issues, we hypothesised methods to increase students’ independent access to research, allowing them to design and undertake useful experiments from first principles. Here, we describe the methods allowing a group of undergraduates to successfully devise and completed a research project independent from their university and academic functions.

## Introduction

Practical research performed by undergraduate students has long been an educational approach to research, primarily useful for the students undertaking a research project. The typical manner of pairing one or a few students with a senior researcher acts as a unique conduit for new students to learn a discipline, and potentially contribute to the field.

Here, we speculate a different approach to undergraduate research, supported by a year long trial involving university students and staff members. We tested the feasibility of performing undergraduate research independent from any existing project or senior research staff. The theory, hypothesis, funding, methods, lab work and dissemination of results was all performed by the same group of undergraduates used during the project.

It is hypothesised that performing undergraduate research from first principles, separated from existing expertise at university, will promote the inquisition, basic skills and care for lab work, and administrative considerations relevant for research in students. The aim is to improve students’ base understanding of research methods and outcomes before embarking on a traditional undergraduate research scheme. A more experienced undergraduate would enhance their learning outcomes of a partnership with an existing project or researcher, as well as having a better ability to contribute useful, novel information to the field.

The specific project designed begins to look at a new form of genetic diagnostic test. This project was designed to run for nine months, giving undergraduate students the opportunity to design, direct and perform experiments to begin characterising key information about this novel test. The project was supported totally internally, using lab facilities and funding made available via internal sources. This emulates the timeline of a more traditional undergraduate research placement. However, the project design, aims and execution are all performed by undergraduates.

## Methods

### Project considerations

The initial project to be undertaken by these undergraduates was designed with some key considerations (CUR, 2024). It must concern recent or breakthrough science, have realistic avenues for improvement and experimentation, and involve limited specialist equipment.

Projects concerning recent science would give undergraduate students relevant experience that may be translated into novel contributions to a related field. Realistic avenues for improvement would allow clear project goals to be defined, generating a direction that would be essential for researchers with limited experience. Limited specialist equipment ensures the project may actually be pursued by undergraduates, without too many funding or safety roadblocks. As such, the scientific methods described herein aim to meet these three key considerations for independent undergraduate research.

### Logistics: finding a laboratory

To get a group of undergraduates performing their own research project requires a laboratory space that can provide the necessary equipment, supervision, storage and safety requirements for biological research. Such a space is often at a premium and presents the biggest roadblock to this kind of work.

Universities that offer lab space to students for extracurricular projects are well positioned to take on this kind of student project. Imperial College is uniquely placed in offering the Advanced Hackspace, containing biolab equipment and supervision. The Hackspace offers a unique sandbox to develop undergraduate research. After approaching Hackspace staff, they receptively opened the space during weekday evenings. This agreement allowed groups of around 20 undergraduate students to work on the project, starting at the beginning of the academic year (October).

Finding a suitable location for undergraduates to complete independent research appears to be one of the key limitations to this kind of pedagogical approach. Such spaces are sometimes available for students to pursue extracurricular biological research, such as what the Hackspace offer. However, they are not routinely used as a supplement to undergraduate teaching. The Hackspace typically excels in supporting small groups of students, e.g. a pair, working towards an idea that may be developed into a product. Such projects are typically funded by winning competition grants, both internal and external to the college.

### Funding

Undergraduates performing lab-based research independently from their university is unusual, and as such, funding presents a large barrier to entry. It was important to keep any funding streams as simple and sustainable as possible. Two options were thus presented.

Firstly, projects may be designed that pursue a new, product-driven line of biological research. Such projects may then be suggested for internal or external funding competitions, such as the Venture Catalyst Challenge offered at Imperial College’s Enterprise Lab. However, competition funding is limited for this idea as caps are often imposed on student numbers, and project funding is often short term.

Alternatively, funding may be pursued internally in the interest of student enrichment. Such funding pools are typically available for student clubs, societies and projects with no immediate link to any academic syllabus. Internal funding from a student union body would be ideal, however difficult, mainly because a project of this kind is linked to the academic development of students, rather than social extracurricular enrichment.

To provide the capital for this project, a balance was struck between these two sources of funding. The project was designed such that potential future outcomes may be commercialised, and was entered into a funding competition with other projects (mostly me chanical e ngine e ring base d) at the Hackspace. The Department of Life Sciences at Imperial College was approached as a source of internal funding. After discussing the potential impact for students, £500 was negotiated to pilot this idea.

The £1000 in funding was used to buy reagents in a quantity large enough to support multiple undergraduates. This funding managed to get an average of five undergraduate students into the lab for 2 hours a week, for 39 academic weeks. Numerically, the funding was aiming to support 400 hours of contact time in the lab, working on the project.

### Project design

Relatively recent methods for isothermal amplification of DNA fragments have been suggested as methods for identifying pathogenic bacteria in foodstuffs (Sun, X. et.al, 2020). Here, DNA fragments matching pathogenic sequences are isolated and identified using pre-made complimentary DNA oligos, rather than thermocycling as in a PCR reaction. The complimentary oligos are fluorescently labeled, and attached to a metal-organic framework (MOF). The MOF quenches fluorescence until such complimentary DNA is available to desorb the oligos and release the fluorophore, identifying the presence of pathogenic sequences of DNA (Boonbanjong, P. et.al, 2022).

This technique uses the CRISPR-Cas9 system (Jinek, M. et.al, 2012) to cut target DNA, and utilises interesting forms of DNA amplification to generate a suitable quantity of target DNA to bind and displace the fluorescent oligos. Isothermal DNA recognition clearly has potential uses as a diagnostic alternative to PCR, if the technology could be made transportable and reliable. This area of research thus satisfies the breakthrough science requirement for such an undergraduate project.

Isothermal amplification methods currently require precise lab conditions to work. These results give no indication to the stability of a MOF carrying fluorescent oligos, or the ability of such a system to work in field conditions where a diagnostic test would be most useful (i.e. lack of temperature control, shelf life etc.). As such, there is a realistic avenue for improvement to this work by identifying such metrics about elements of the system.

Overall, the aim of the project was twofold. First, to explore the synthesis of a relevant MOF using rudimentary lab equipment and expertise. Second, to explore the ability of such a MOF to bind to and quench fluorescence of labelled DNA oligos. This would give an interesting insight to the robustness of such a system to one-pot, ‘field like’ conditions that a useful diagnostic test using this technology might encounter. This focus also removes any requirements for live biologicals (bacterial cultures) to be handled, removing a potential safety roadblock to the research.

### Scientific methods

The MOF to be synthesised was UiO66 (Cavka, J.H. et.al, 2008). To produce the UiO66, reagents were added to a 100mL teflon-lined steel autoclave. 19mg of Zirconium (IV) Chloride and 14mg Terepthalic acid (TA) were weighed, and added to 2mL Acetic Chloride and 10mL Dimethylformamide (DMF), directly to the teflon carrier. This solution was ultrasonicated for 30 minutes, followed by an 8 hour autoclave cycle at 120C (Sun X. et.al, 2020).

The resulting product was collected from the teflon carrier into 50mL Falcon tubes. The tubes were spun at 4000 RPM for 20 minutes, and solids separated. The solids were left overnight in a vacuum desiccator to dry. These dried solids were expected to be the UiO66. This solid was then used to assess DNA binding.

To assess binding, varying amounts of the solid MOF were dissolved in Tris buffer. The fluorescently labelled DNA oligos were added to these solutions, and emission spectra used to calibrate the plate reader to the correct excitation range. Peak absorbance was measured at 470nm, the output wavelength of the FAM-label (Gesner, T., Mayer, U. 2000).

The scientific methods were designed to be robust, yet vague. In brief, the method was to mix reagents, produce a solid after an autoclave cycle, separate and dry that solid, before mixing it with fluorescent DNA. Each preparation thus had a certain variance in amounts of reagents used, time spinning down solids and the drying time/dry weight.

## Results

The results discussed are in two parts. The first considers the student outcomes in completing this project, and the potential outcomes of future repeats of a similar method. The second considers the scientific outcome of the results.

### Student outcomes

It is difficult to quantitate the impact such research experience has on students unless their academic outcomes can be accurately compared to a control group, over a longer term. Short term metrics can be discussed however. A simple cost-benefit analysis may be performed by comparing the money spent on the project against the number of repeat attending students.

£1000 was spent in total on this project. The numbers predicted by our methods remained largely true, supporting 5 to 10 undergraduates a week for almost the entire academic year (39 weeks), for two hours a week. At the top end of attendance, this project could provide upwards of 1,100 contact hours for students, in the lab, working on a project they have designed.

Some qualitative reviews from undergraduates about their experience of this project are available in the appendix.

### Scientific results

The scientific results were twofold. Firstly, the MOF was synthesised. Secondly, its ability to bind fluorescently labelled DNA probes and quench fluorescence was tested. After the synthesis protocol was followed, the MOF was imaged using scanning electron microscopy within the Department of Chemical Engineering (fig.2).

**Figure 1:**
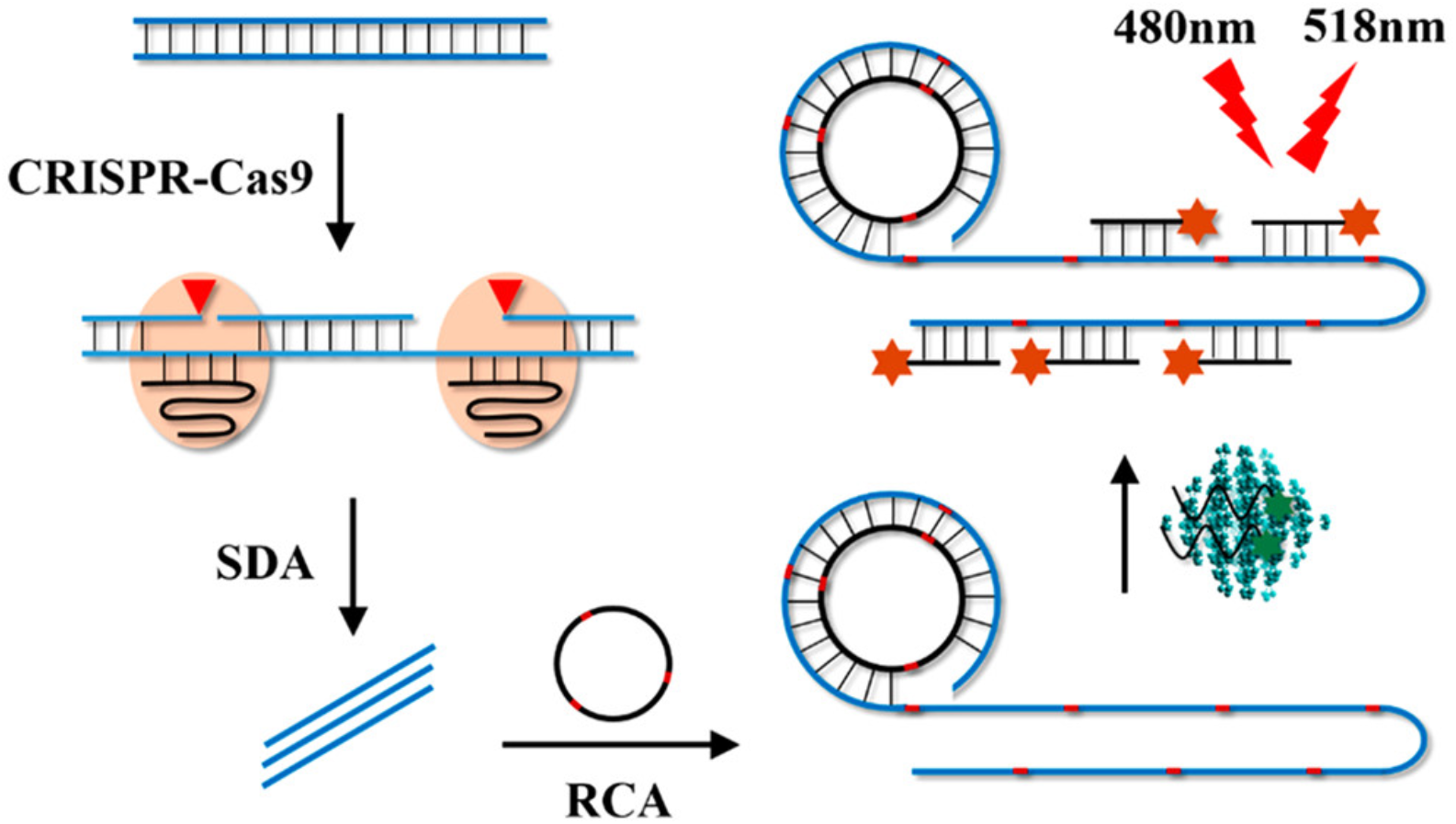
Schematic of the method to detect specific genetic material by isothermal amplification. Taken from Sun, X. et.al, 2020.

**Figure 2.**
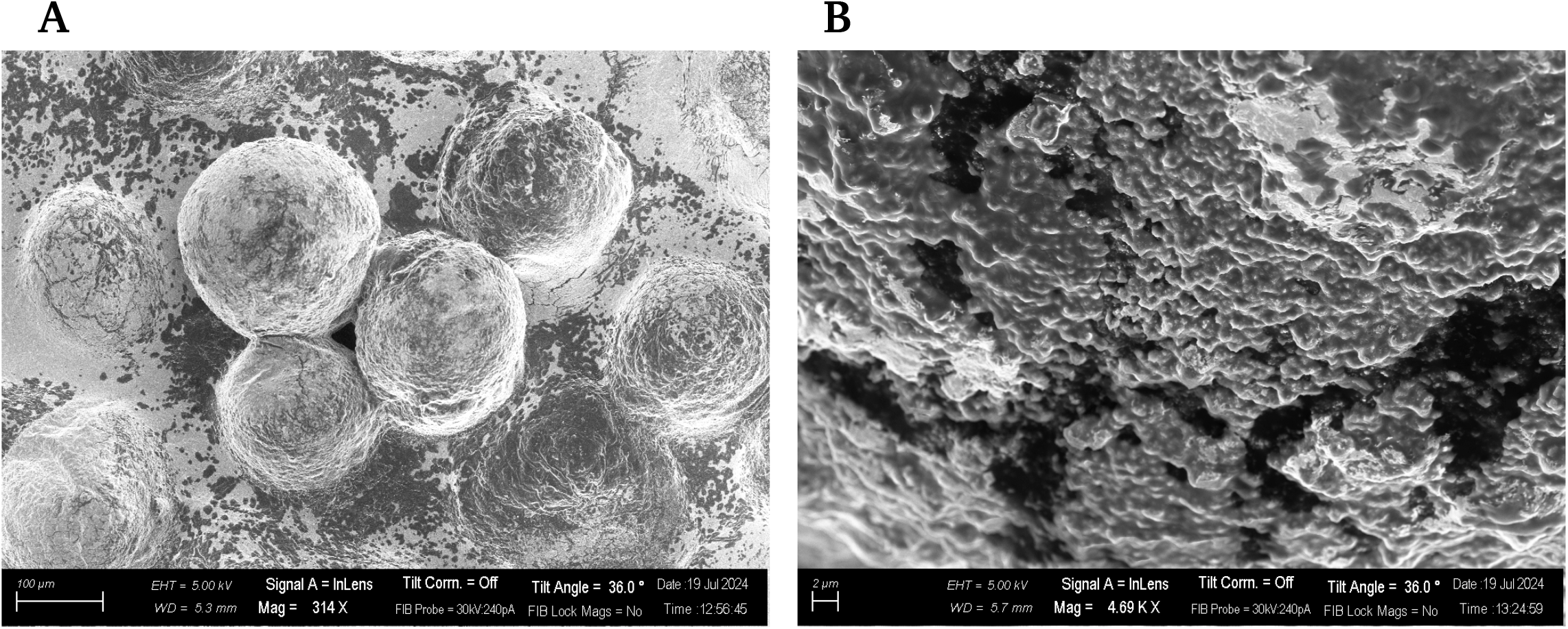
**(A)** 100μm scale SEM image of a sample of UiO66 produced by undergraduates. The pea-like structures suggest a repeating crystal lattice, solvated by reagents not fully removed during the SEM preparation process. **(B)** A deeper, 2μm scale SEM image of a sample of UiO66 produced by undergraduates. The rough surface suggests a crystal lattice was obtained, as expected from other samples of UiO66 produced using a similar method.

The images suggest a form of crystalline structure was obtained, due to the repeating unit nature of the sample. Further, some of the deeper images imply geometry similar to that observed in other UiO66 images. Clearly, the sample tested did not display neat and clear crystal units. The sample appears to be solvated, most likely stemming from residual synthesis reagents not completely removed during SEM preparation.

Only one sample was able to be imaged due to time constraints in accessing the SEM. Therefore, it is hard to say if this morphology is representative of all the UiO66 produced during the project. However, having an image of what was produced was incredibly valuable for project morale and as a project deliverable.

The UiO66 sample imaged in the SEM was the same used to test DNA binding. Here, a plate reader was used to measure the fluorescence of the FAM labelled DNA probes in the presence of differing concentrations of UiO66 (following appendix table).

Figure 3 shows a significant trend between absorbance of DNA solutions. At 12.5% UiO66 concentration (a suggested maximum amount from Sun, X.), absorbance is low, suggesting fluorescent DNA has been quenched by UiO66. As concentration decreases, more DNA is available to fluoresce, and absorbance increases.

**Figure 3:**
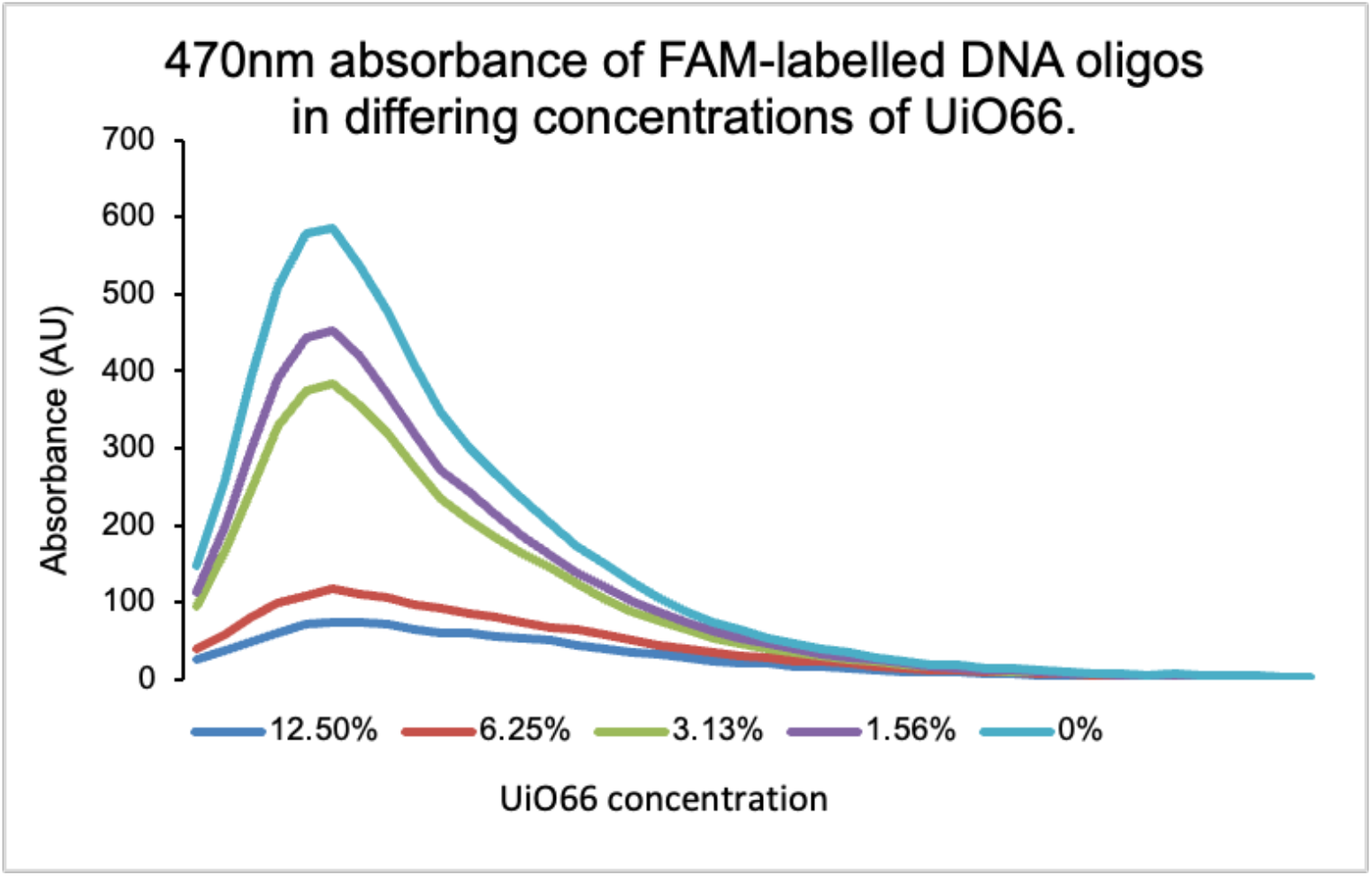
emission spectra of fluorescently labelled DNA oligos. Oligos were prepared in the solvents used for UiO66 synthesis. Samples were excited at 470nm, and emission measured between 495-700nm.

This trend was controlled by measuring fluorescence of different concentrations of FAM-oligos independently, and identifying the trend. This is effectively how UiO66 affects fluorescence, by changing the concentration of fluorophore available in solution.

## Discussion

This project aimed to describe an approach to get undergraduate Life Science students completing their own, laboratory research given the particular facilities at a particular university. The project has resulted in some interesting scientific contributions to the field of metal organic framework diagnostics, and a pedagogical discussion about undergraduate research in the Life Sciences.

We find the cost-benefit ratio of funding such a project has potential to be very high. Certain institutions will be better placed along this ratio, such as Imperial College, with their Advanced Hackspace offering highly accessible lab space.

This project aimed to produce a relatively new inorganic crystal, UiO66. The potential for this crystal to bind and quench fluorescence of labelled DNA oligos was the main question during this project. Such properties lend similar crystals for use in isothermal diagnostic tests, able to recognise specific DNA sequences via fluorescence.

Our results suggest the ability of UiO66 to quench fluorescence when provided with labelled DNA oligos. This was expected, and shows the ability of undergraduate research to faithfully reproduce experiments. What was unexpected was the level of acceptable variability when producing and storing UiO66.

This project was able to demonstrate the isothermal, functional fluorescence quenching across a range of circumstances relevant to field use diagnostics. The fact this was done entirely by undergraduates, with limited funding, demonstrates the usefulness and need for this kind of undergraduate research, for students and research fields more generally.

*J*.*G hypothesised the research question, developed project logistics and practical experiments, and compiled the manuscript. A*.*B. helped develop the project logistics, aims, and review of the manuscript. H*.*B. provided access, training and guidance for SEM images*.

## Appendix

### Student testimonials

#### Basem Qubaty. 2nd year, Biochemistry

My experience in the lab has been great so far. I have been able to get some hands on experience with many procedures; this has helped me to solidify my understanding of theoretical principles that I have encountered during my course. The program has certainly benefited my academic development and has made me more confident with carrying out lab procedures. I look forward to gaining even more valuable practical experience during the next stage of the program.

#### Kai Mulcock. 4th year, Physics

As the first bit of biological lab work I have done it was a great introduction! It was helpful having fellow students lead the class as I felt a lot more comfortable and confident in asking and learning about different topics (despite not knowing any biological background beyond A level).

As I am not a biology student I cannot speak too much on the theories, however, it was very interesting seeing application of even some things from Physics!

It provided a useful and necessary step in helping me know if this is a field I would like to explore more, while also showing me what niches I could fill as a non biological student!

#### Jack White. 3rd year, Materials Science

My experience in the lab has been incredibly rewarding and enlightening. Working hands-on with biological materials has allowed me to apply theoretical knowledge from my materials science background in a practical context.

The collaboration with biologists and exp osure to advanc ed tec hni q ues have broadened my perspective and sparked my interest in the intersection of materials science and biology. If I were speaking to department leaders, I would emphasize that this program has significantly impacted my academic and personal development. Academically, it has enhanced my research skills, critical thinking, and problem-solving abilities.

The experience has cultivated my ability to work collaboratively in interdisciplinary teams, which is crucial in today’s research environment. Personally, it has boosted my confidence in conducting scientific research and ignited a passion for exploring how materials can be engineered for medical and biological applications.

**Appendix table:** amounts of reagents added across three rows of a 96-well plate used for assessing fluorescence. MOF was added last to the relevant wells to help control side reactions with excess Zirconium Chloride used to synthesise the MOF.

|  | 1 | 2 | 3 | 4 | 5 | 6 |
| --- | --- | --- | --- | --- | --- | --- |
| A | ⇒100μL Tris | ⇒100μL Tris | ⇒100μL Tris |  |  |  |
| B | ⇒12.5μL MOF<br>⇒50μL FAM-Oligo<br>⇒37.5μL Tris | ⇒6.25μL MOF<br>⇒50μL FAM-Oligo<br>⇒43.75μL Tris | ⇒3.125μL MOF<br>⇒50μL FAM-Oligo<br>⇒46.875μL Tris | ⇒1.56μL MOF<br>⇒50μL FAM-Oligo<br>⇒48.44μL Tris | ⇒50μL Tris<br>⇒50μL FAM-Oligo |  |
| C | ⇒100μL FAM-Oligo | ⇒80μL FAM-Oligo<br>⇒20μL Tris | ⇒60μL FAM-Oligo<br>⇒20μL Tris<br>⇒20μL Water | ⇒40μL FAM-Oligo<br>⇒20μL Tris<br>⇒40μL Water | ⇒20μL FAM-Oligo<br>⇒20μL Tris<br>⇒60μL Water | ⇒20μL Tris<br>⇒80μL Water |

#### Appendix table

Volumes of reagents added to each well of a 96-well plate. This plate was used to produce FAM-absorbance figures. An average of row A was used as a blank.

The Tris buffer was used to maintain a slightly acidic pH, and as a medium for fluorescence solutions. It would be possible to use other buffers that support DNA and the MOF crystal (namely, a strong neutral pH buffer). Tris was chosen for its availability, cost and robustness.

**Appendix figure:**
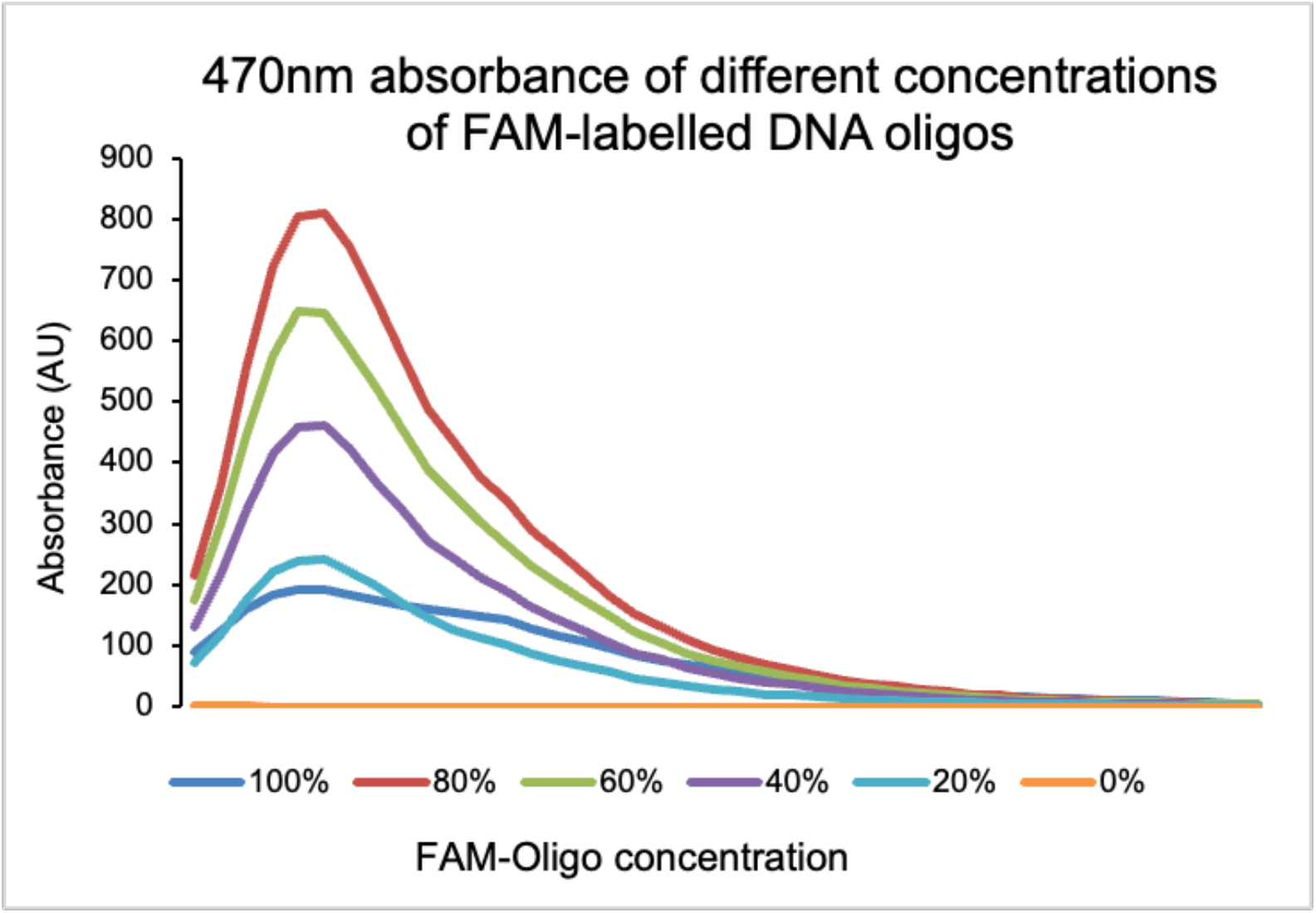
control fluorescence readings, plotted using appendix table 1, row C. There is a slight anomaly at a concentration of 100%, however the trend upholds.

## Notes

### Competing Interest Statement

The authors have declared no competing interest.

